# Temperature modulates PVN pre-sympathetic neurones via transient receptor potential ion channels

**DOI:** 10.1101/2022.01.26.477880

**Authors:** Fiona O’Brien, Claire Feetham, Caroline Staunton, Katharine Hext, Richard Barrett-Jolley

## Abstract

**Background and purpose:** The paraventricular nucleus (PVN) of the hypothalamus plays a vital role in maintaining homeostasis and controls cardiovascular function via autonomic pre-sympathetic neurones. We have previously shown that coupling between transient receptor potential cation channel subfamily V member 4 (Trpv4) and small-conductance calcium-activated potassium channels (SK) in the PVN facilitate osmosensing. TRP channels are also thermosensitive and therefore, in this report we investigated the temperature sensitivity of PVN neurones.

**Experimental approach:** We identified TRP channel mRNA in mouse PVN using quantitative reverse transcription-PCR (RT-PCR). Using cell-attached patch-clamp electrophysiology, we characterised the thermosensitivity of Trpv4-like ion channels on mouse PVN neurones. Following recovery of temperature sensitive single channel kinetic schema, we constructed a novel and predictive stochastic mathematical model of these neurones. We then validated this model with electrophysiological recordings of action current frequency from mouse PVN neurones.

**Results:** We identified 7 TRP channel genes in the PVN with known thermosensitive capabilities. Trpv4 was the most abundant of these and was easily identified at the single channel level using cell-attached patch-clamp electrophysiology on PVN neurones. We investigated the thermosensitivity of these Trpv4-like channels; open probability (Po) markedly decreased when temperature was decreased, mediated by a decrease in mean open dwell times. Our neuronal model predicted that PVN spontaneous action current frequency (ACf) would increase as temperature is decreased and in our electrophysiological experiments, we found that ACf from PVN neurones was significantly higher at lower temperatures. The broad-spectrum channel blocker, gadolinium (100 μM), was used to block the warm-activated Ca^2+^-permeable Trpv4 and Trpv3 channels. In the presence of gadolinium (100 μM), the temperature effect was largely retained. Using econazole (10 μM), a blocker of Trpm2, we found there were significant increases in overall ACf and the temperature effect was inhibited.

**Conclusion:** Our work identified Trpv4 mRNA as an abundantly expressed thermosensitive TRP channel gene in the PVN and this ion channel contributes to the intrinsic thermosensitive properties of PVN neurones. At physiological temperatures (37°C), we observed relatively low ACf primarily due to the activity of Trpm2 channels, whereas at room temperature, where most of the previous characterisation of PVN neuronal activity has been performed, ACf is much higher, and appears to be predominately due to reduced Trpv4 activity. This work gives insight into the fundamental mechanisms by which the body decodes temperature signals and maintains homeostasis.

## Introduction

Prolonged deviation in core body temperature (T_c_) outside a narrow range results in serious physiological issues incompatible with life, and therefore, it is tightly regulated by a homeostatic system (Gomez, 2014; Morrison & Nakamura, 2011). Changes in environmental temperature produce reflex responses to maintain T_c_ in an optimal range (Kanosue et al., 1991; Kanosue, Yanase-Fujiwara & Hosono, 1994), however, how the brain coordinates such responses is a longstanding and unresolved question.

Early models of temperature regulation were based around the existence of a central integrator comprised of hypothalamic neurones that orchestrate homeostatic responses around a set-point temperature (H T Hammel & Pierce, 1968; Hardy, 1961). An alternative theory proposes that the brain has no central integrator for T_c_ but instead, various thermoregulatory effectors are thought to be regulated independently, giving the appearance of coordinated action without the existence of a single so-called ‘controller’ (McAllen, Tanaka, Ootsuka & McKinley, 2010; Romanovsky, 2007a; Satinoff, 1978). Central nervous system (CNS) level control of T_c_ is mediated by a combination of negative feedback and feed-forward mechanisms that share common peripheral thermal sensory inputs (Kanosue, Crawshaw, Nagashima & Yoda, 2010; Morrison & Nakamura, 2011). Feedback responses are those that are triggered when T_c_ deviates away from the optimal range, for example, exercise induces an increase in T_c_ by several degrees Celsius (Fuller, Carter & Mitchell, 1998; Walters, Ryan, Tate & Mason, 2000). Feed-forward mechanisms on the other hand, are preventative, and are triggered prior to any change in core temperature. The most common feed-forward example is the detection of change in air temperature (by thermoreceptors in the skin) which trigger thermoregulatory responses that prevent any significant change in T_c_ (Nakamura & Morrison, 2008; Nakamura & Morrison, 2010; Romanovsky, 2014). The hypothalamus contains the primary integrative and rostral efferent components of these circuits, but local thermal stimulation of other areas in the CNS, including several brain-stem neuronal groups and the spinal cord also trigger autonomic thermoeffector responses. Thermosensitive neurons in the preoptic area (POA) of the hypothalamus have been the most studied to date (Kanosue et al., 1991; Kanosue, Yanase-Fujiwara & Hosono, 1994; Kazuyuki, Hosono, Zhang & Chen, 1998; Nagashima, Nakai, Tanaka & Kanosue, 2000; Nakamura & Morrison, 2010; Romanovsky, 2007b).

Early experiments showed that stimulation (warming) of the cat (Hemingway, Forgrave & Birzis, 1954; Magoun, Harrison, Brobeck & Ranson, 1938) and rat (Carlisle & Laudenslager, 1979) POA could trigger dramatic thermoregulatory responses that were similar to those observed by heating the entire animal. Cooling of the POA promotes vasoconstriction, BAT thermogenesis and shivering in dogs and baboons (Gale, Mathews & Young, 1970; Hammel, Hardy & Fusco, 1960) and results in baboons signalling for rapid heat reinforcement. Lesioning of the cat POA has been shown to abolish thermoregulatory responses in animals subjected to temperature challenge (Clark, Magoun & Ranson, 1939; Teague & Ranson, 1936). Direct sensing of changes in skin temperature has been shown to activate POA efferent signals that control thermal effector organs (Morrison, 2016; Morrison, Madden & Tupone, 2014). Electrophysiological studies have characterised the intrinsic temperature-sensitive properties of POA neurones in rabbit (Boulant & Hardy, 1974), rat (Baldino & Geller, 1982; Hori, Nakashima, Hori & Kiyohara, 1980) mice (Tan et al., 2016) and dogs (Hardy, Hellon & Sutherland, 1964). However, there have also been many reports of temperature sensitive neurones outside of the POA (Edinger & Eisenman, 1970; Kobayashi & Murakami, 1982; Nakayama & Hardy, 1969; Simon & Iriki, 1970; Wünnenberg & Hardy, 1972). The neuronal circuitry and projections of the POA are not fully understood but several additional brain regions including the dorsomedial hypothalamus (DMH), the paraventricular nucleus of the hypothalamus (PVN), and the raphe pallidus nucleus have been proposed to act alongside the POA to regulate T_c_ (Morrison, 2016; Morrison, Madden & Tupone, 2014; Zhao et al., 2017). The DMH is also recognised as another key player in thermoregulation (Dimicco & Zaretsky, 2007; Heeren & Münzberg, 2013; Morrison & Nakamura, 2011) and stimulation of rat DMH neurons was shown to increase in BAT sympathetic nerve activity (SNA), BAT temperature and TC (Cao, Fan & Morrison, 2004; de Menezes, Zaretsky, Fontes & DiMicco, 2006; Zaretskaia, Zaretsky, Shekhar & DiMicco, 2002).

Several studies using cFos as a marker of activation have shown that mouse PVN neurones respond to both warm and cold ambient temperature change (Bachtell, Tsivkovskaia & Ryabinin, 2003; Bratincsák & Palkovits, 2004). Exposure to a hot environment (39°C) increased cFos expression of rostral ventrolateral medulla (RVLM) –projecting (Cham & Badoer, 2008) and spinally-projecting neurones in the rat PVN (Cham, Klein, Owens, Mathai, McKinley & Badoer, 2006). Anatomical studies using transneuronal viral tracing approaches show that post injection of pseudorabies virus into the rat tail, within the hypothalamic area, the majority of labelled neurons were located in the PVN (Smith, Jansen, Gilbey & Loewy, 1998). Injection of glutamate in the PVN leads to an increase in interscapular brown adipose tissue (IBAT) temperature in rats and on the other hand, lesioning of the PVN reduced febrile-evoked increases in body temperature, suggesting a role for the PVN in driving sympathetic outflow to BAT, at least in the context of fever (Amir, 1990; Caldeira, Franci & Pelá, 1998; Horn, Wilkinson, Landgraf & Pittman, 1994; Leite, Zheng, Coimbra & Patel, 2012; Lu et al., 2001). Furthermore, more generally, Cabral *et al.*, (2012) showed that TRH (a neuropeptide necessary for cold-induced thermogenesis) -neurones in the rat PVN are activated when animals are exposed to short-term cold conditions (Cabral, Valdivia, Reynaldo, Cyr, Nillni & Perello, 2012).

Electrophysiological studies have been pivotal to not only characterising the biophysical profile of PVN neurones, but also understanding how the PVN plays a role in the regulation of homeostatic functions. The PVN is typically divided into the parvocellular and magnocellular regions, which both have many subdivisions (BÜTtner-Ennever, 1997; Koutcherov, Mai, Ashwell & Paxinos, 2000) but in its most simplistic representation, the PVN is divided loosely into the parvocellular area, posterior magnocellular lateral area and the intermediocellular region (dorsal and PVN) where pre-autonomic neurones are most abundant (Chen, Gomez-Sanchez, Penman, May & Gomez-Sanchez, 2014; Feetham, O’Brien & Barrett-Jolley, 2018; Kiss, Martos & Palkovits, 1991). To date, three distinct electrophysiological phenotypes have been described in PVN neurones (reviewed in Feetham *et al.*, (2018)); magnocellular (type I) PVN neurones have phasic bursting patterns and express a rapidly inactivated, or “A-type,” potassium conductance (Sonner & Stern, 2007; Tasker & Dudek, 1991). Parvocellular (type II) PVN neurones express a slowly inactivating delayed rectifier potassium conductance and it is suggested that the differences between types I and II cells may be explained by differential expression of voltage-gated potassium and calcium channels (Luther, Daftary, Boudaba, Gould, Halmos & Tasker, 2002). Furthermore, in the parvocellular area, there appears to be two different neuronal phenotypes; (1) exhibiting electrophysiological properties similar to neuroendocrine magnocellular cells (Stern, 2001; Tasker & Dudek, 1993) and (2) pre-autonomic/spinally projecting neurones which show neurones a slowly inactivating potassium conductance (Barrett-Jolley, Pyner & Coote, 2000; Tasker & Dudek, 1993). In regards to thermosensitivity, Inenaga *et al.*, (1987) was the first study to confirm the inherent thermosensitivity of PVN neurones and characterised separate intrinsically “cold-sensitive” and “warm-sensitive” neurones (Inenaga, Osaka & Yamashita, 1987). To our knowledge, this is the only electrophysiological study investigating the temperature sensitivity of PVN neurones, and to date, there has been no molecular characterisation.

The cellular pathways involved in thermo-sensation are well conserved and consist of a set of specialised temperature-gated ion channels that are highly sensitive to a wide range of hot and cold temperatures. The thermo-transient receptor potentials (TRPs), a recently discovered family of ion channels activated by temperature, are expressed in primary sensory nerve terminals where they provide information about thermal changes in the environment. There are 4 heat thermo-sensitive TRP ion channels; Trpv1 (activated with temperature >43°C) (Everaerts et al., 2011), Trpv2 (activated with temperature >52°C) (Liu & Qin, 2016), Trpv3 (activated with temperature >32°C) (Peier et al., 2002) and Trpv4 (activated with temperature >27°C) (Güler, Lee, Iida, Shimizu, Tominaga & Caterina, 2002). In addition, there are 2 identified cold thermo-sensitive TRP ion channels; Trpm8 (activated with temperature <28°C) (McKemy, Neuhausser & Julius, 2002) and Trpa1 (activated with temperature <17°C) (Laursen, Bagriantsev & Gracheva, 2014). PVN ion channels, including those that are thermo-sensitive have recently been summarised in (Feetham, O’Brien & Barrett-Jolley, 2018).

We have previously shown that Trpv4 is expressed on PVN neurones of CD1 mice (Feetham, Nunn & Barrett-Jolley, 2015; Feetham, Nunn, Lewis, Dart & Barrett-Jolley, 2015). Originally, these channels were considered sensors of cell volume (Liedtke et al., 2000) and in the PVN, we have shown that Trpv4 channels functionally couple to a subtype of Ca^2+^-activated K^+^ channel (SK channel) to sense changes in osmolality, probably mediated by subtle changes in cellular volume (Feetham, Nunn, Lewis, Dart & Barrett-Jolley, 2015). We also found that ICV injection of hypotonic artificial cerebrospinal fluid (ACSF) into CD1 mice decreased mean blood pressure, but not heart rate and this effect was abolished by treatment with the Trpv4 inhibitor RN1734 (Feetham, Nunn & Barrett-Jolley, 2015). In another recent study, we found that systemic administration of the highly selective lipid-soluble Trpv4 antagonist GSK2193874 resulted in tail blood-flow dynamics that were in-compatible with a local (vascular smooth muscle or endothelial cell) mechanism (O’Brien, Staunton & Barrett-Jolley, 2021). Accumulating data shows that Trpv1 may play a role in regulating sympathetic outflow in hyperthermic responses, but in the light of the above data, we hypothesised that PVN Trpv4 ion channels also play a role in thermoregulation (Alawi et al., 2015).

Many studies have shown that Trpv4 can also be activated by heat >25°C (Clapham, 2003; Güler, Lee, Iida, Shimizu, Tominaga & Caterina, 2002; Watanabe, Vriens, Suh, Benham, Droogmans & Nilius, 2002) as well as mechanical stimuli. Trpv4 immunoreactivity is present in a number of brain regions known for producing thermoeffector responses including the POA (Güler, Lee, Iida, Shimizu, Tominaga & Caterina, 2002), the PVN (Feetham, Nunn & Barrett-Jolley, 2015) but also in the vasculature where its activation produces vasodilation (Filosa, Yao & Rath, 2013).

Therefore, the aim of this study was to characterise the thermosensitivity of a subpopulation of PVN neurones; we have already shown that Trpv4 channels are present in the PVN but here, we used RT-PCR to identify additional thermosensitive targets. We pharmacologically identified Trpv4-like channels from PVN neurones and characterised their intrinsic thermosensitive properties at the single-channel level. We built on from our previous mathematical model to predict that neuronal activity should increase as temperature is decreased; we validate this model with recordings from PVN neurones. We show that the temperature-sensing capabilities of PVN neurones is complex, and is likely to involve multiple ion channels, including Trpv4, Trpv3 and another known thermosensitive ion channel, Trpm2.

## Methods

### Animals

Mice were housed at 22-24°C in a 12 h light/dark cycle-controlled facility with ad libitum access to food and water. Animals were sacrificed by Schedule 1 methods for all in vitro work and all experiments were approved by the Home Office, UK.

### Quantitative PCR

Young (6-8 months) and old (24 months) mice were killed by Schedule 1 methods and the general hypothalamic area was blocked and transversely sliced with 600μM thickness. The PVN was identified using the third ventricle and fornix as markers and was punched out using a 1.5 mm biopsy punch. PVN punch biopsies were suspended in 14μL of RNA-free water and homogenised by passing the lysate through 20-gauge needle multiple times. To obtain the required volume of mRNA, samples were pooled, with between 2-3 PVN punch biopsies per 14μL of RNA-free water. RNA extraction was carried out using the RNeasy Plus Micro kit, together with gDNA eliminator and MinElute spin columns (Qiagen, UK) and analysed for concentration, purity and quality using the NanoDrop™ 2000 (Thermo Scientific, UK). cDNA synthesis (mRNA) was performed using the RT2 First Strand kit (Qiagen, NL) according to the manufacturer’s protocol. Using the Neuronal ion channel plate (Qiagen, UK), 84 ion channels as well as housekeepers were measured in each sample. qPCR analysis was performed using the Stratagene MX3000P RT-PCR System (Stratagene, La Jolla, CA) in a 25-μL reaction mixture. Expression relative to mean of 3 housekeeper genes (Actb, Ldha, Rplp1) Cts for that sample is presented as the ΔCt.

### Brain slice preparation

CD1 mice of both sexes, aged 2–3 weeks were killed by Schedule 1 methods and the brain was swiftly removed and placed in ice-cold low Na^+^/high-sucrose artificial cerebrospinal fluid (ACSF) and sliced as previously described (Feetham, Nunn, Lewis, Dart & Barrett-Jolley, 2015). In brief, coronal PVN slices were prepared using a Campden Instruments Ltd Leica VT1000S and stored in a multiwell dish containing physiological ACSF. Slices were kept at 35-37 °C with continuous perfusion of 95% O2/5% CO2 and left to recover for at least 1 hr before recording.

### Electrophysiology

Thick-walled patch-pipettes were fabricated using fire-polished 1.5 mm o.d. borosilicate glass capillary tubes (Sutter Instrument, Novato CA, USA) using a two-step electrode puller (Narishige, Japan). Neurons were visualised using a Hitachi KP-M3E/K CCD camera attached to a Nikon Eclipse microscope with an effective magnification of ~1000x. Cell-attached patch clamp electrophysiology was performed as previously described using an Axopatch 200b amplifier (Molecular Devices Axon Instruments, USA) (Feetham, Nunn, Lewis, Dart & Barrett-Jolley, 2015). For spontaneous action current recordings, analogue data were further amplified with a Tektronix FM122 (Beaverton, OR, USA) AC-coupled amplifier. The temperature of the recording bath was maintained using the npi electronic TC-10 (Scientifica, UK). In all cases, data were low-pass filtered at 1 kHz and digitised at 5 kHz with a Digi Data 1200B interface. Recording solutions are described below and junction potentials were calculated using JpCalc (Barry & Lynch, 1991).

### Analysis of electrophysiological recordings

Single channel recordings were digitally filtered at 1 kHz in WinEDR (University of Strathclyde, UK). Open and closed levels were assessed by all-points amplitude histograms and were used to create current-voltage (IV) curves. Single channel events were idealized using the segmental K means (SKM) methods (Qin, 2004) using QuB software (SUNY, Buffalo, NY) and open probability (Po) was determined from the idealised record as previously described (Lewis et al., 2013). For dwell time analysis, dead-time was set to three sample intervals (0.3 ms) and recordings where only a single channel was gating were used. Open and closed dwell times were log binned according to the methods of (Sigworth & Sine, 1987) and fitted with an exponential log probability density function (pdf) in Clampfit 10.3 (Molecular Devices, Sunnydale California). The number of time constants for each distribution was determined using a log-likelihood ratio test in Clampfit 10.3 at a confidence level of P = 0.95.

Individual sets of model kinetic rates were obtained by fitting the idealised data using the MIL algorithm implemented in QuB. Missed events during maximum interval likelihood (MIL) rate optimisation were automatically accounted for in QuB by computation of a corrected Q matrix in the MIL algorithm.

Analysis of action current frequency was performed using WinEDR for acquisition of data and then a custom program designed to detect action currents based on an adaptive threshold routine.

### Solutions

Cell-attached patch recordings were made using the following solutions: ACSF composition (mM): 127 NaCl, 1.8 KCl, 1.2 KH_2_PO4, 2.4 CaCl_2_, 1.3 MgSO_4_, 26 NaHCO_3_ and 5 glucose. Pipette solution for action current and single channel recordings composition (mM): 35 KG; 5 KCl; 100 NaCl and 10 HEPES (pH 7.4) with NaOH. All experiments were performed in the daytime (11:00–17:00 h) to limit the effects of circadian rhythm on activity of the cells used (Belle, Diekman, Forger & Piggins, 2009).

### Design of the computer model

Mathematical models were constructed in Python using open-source NEURON libraries (Hines & Carnevale, 1997a; Hines, Davison & Muller, 2009). Our model was based on that of Feetham *et al.*, 2015 (Feetham, Nunn, Lewis, Dart & Barrett-Jolley, 2015); in brief, inputs arise from both excitatory ‘Netstim’ neurones and inhibitory interneurons, see Fig. 5A. The interneurones are also driven by excitatory ‘Netstim’ neurones. Since computer power has increased significantly, the model has been updated in several ways: (a) The new model uses stochastic channels rather than “density” (deterministic equations), since the noise added by stochastic simulation allows for more authentic simulation (Cannon, O’Donnell & Nolan, 2010). To obtain single channel rate constants for Kv channels we fitted our whole-cell Kv data using a Monte Carlo bootstrap approach within Python. Netstim activities are also now scattered stochastically around the fixed means previously used by Feetham *et al.*, 2015 (Feetham, Nunn, Lewis, Dart & Barrett-Jolley, 2015). (b) We replaced osmotic sensitive Trp4 with temperature sensitive channels, using our experimentally measured rate constants, stochastic model and conductance, see (Fig. 5A). Temperature dependence was included by applying Q10 to each of the forward rate constants. We also added stochastic SK channels from the model of Moczydlowski and Latorre (Moczydlowski & Latorre, 1983) and a hypothetical TRPM2-like channel using arbitrary base rate-constants, but Q10 (15.6) measured by Togashi *et al.*, 2006 (Togashi et al., 2006). Ion channel permeabilities were from Alexander *et al.*, 2015 (Alexander et al., 2015). (c) We replaced the former bulk Ca^2+^ accumulation mechanism (Feetham, Nunn, Lewis, Dart & Barrett-Jolley, 2015) with a new reaction diffusion (RXD) model (McDougal, Hines & Lytton, 2013b) including central Ca^2+^ ion buffering. All code will be made freely available on GitHub, and if possible, ModelDB.

### Data analysis

All data on graphs are shown as mean ± SEM. Simple comparisons were made using a two-tailed Student’s paired t-test. Multiple comparisons were made using a repeated measures ANOVA with multiple comparisons by Tukey’s post hoc test or against control levels using Dunnett’s post hoc test where appropriate. A value of P < 0.05 was taken as significant.

### Materials

GSK2193874 (80 nM), gadolinium (100⍰μM) and Econozole (10⍰μM) were sourced from Sigma-Aldrich and were all dissolved in DMSO and diluted to a final working concentration of no more than 0.01% DMSO (0.01% DMSO had no effect alone).

## Results

### Gene expression levels of thermosensitive TRP channels in punches of mouse PVN

We measured mRNA (by quantitative qPCR) of the warm activated Tprv1 (DCt 11.50±2.97, n=3), Tprv2 (DCt 5.29±0.32, n=3), Tprv3 (DCt 9.39±0.35, n=3), Tprv4 (DCt 4.46±0.22, n=3) and Trpm2 (DCt 4.04±0.36, n=3) channels and mRNA levels of the cold activated Trpm8 (DCt 8.78±1.50, n=3) and Trpa1 (DCt 8.48±2.34, n=3) channels (Klein, Trannyguen, Joe, Iodi & Carstens, 2015; Nazıroğlu & Braidy, 2017). Note lower DCt means higher mRNA abundance. Therefore, in young mice (the age used in the rest of this work) Trpv4 and Trpm2 were the most abundantly expressed of these thermosensitive TRP channels (Fig. 1). Full datasets for a set of 84 ion channel genes including these, in both young and old mice are included in the supplementary information.

**Figure 1.**
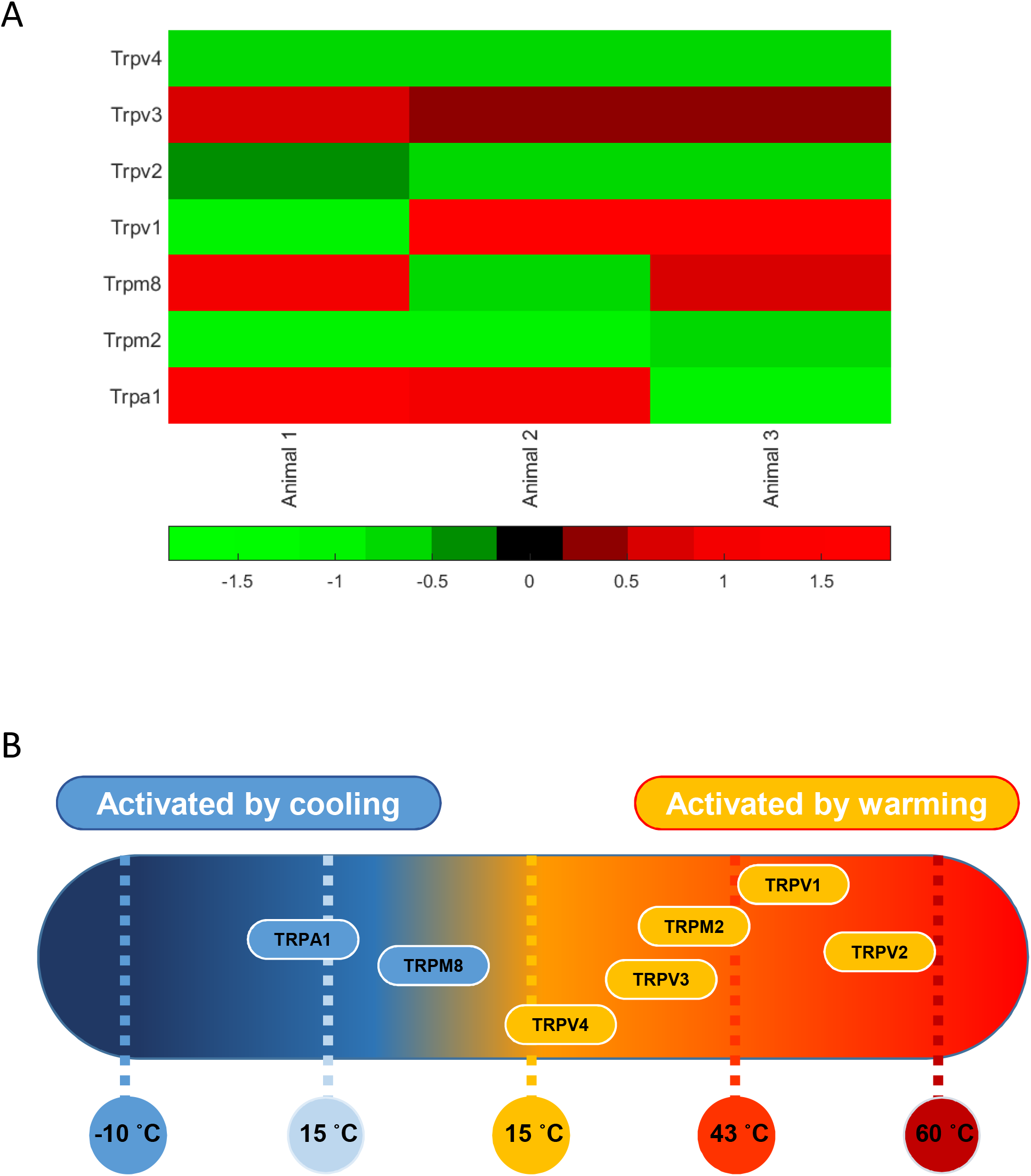
Thermosensitive TRP channel gene expression in PVN punches from young mice. **A.** Standardized DCt levels for seven known TRP channel mRNA in punches of the PVN (from 3 animals). Since this DCt, the green (negative values) are relatively high expression and the red (positive values) are low expression. Mean DCt was lowest (highest mRNA abundance) for TRPV4 (see values in the text). A full dataset of 84 ion channel mRNA DCt levels measured in younger (6-8 month) and older (26 month) mice are given together with heatmaps in the supplementary information. **B.** Established distribution of transient receptor potential (TRP) channels in PVN tissue as a function of their temperature threshold. TRP channels may be activated by increases in temperature (orange) or by lowering the temperature (blue) (Klein, Trannyguen, Joe, Iodi & Carstens, 2015). Image modified under BY4.0 Creative Commons licence from (Lamas, Rueda-Ruzafa & Herrera-Pérez, 2019).

### Identification of Trpv4 channels on PVN neurones

To identify and characterise the single-channel gating of PVN Trpv4 channels, we used cell attached electrophysiology on PVN neurones. A Trpv4-like channel was identified in 56% of recordings with a conductance of 59.7 ± 1 pS and Vrev of −5.9 ± 3 mV (Fig. 2A(i), n=10). This channel was absent when cells were patched in the presence of the specific Trpv4 antagonist GSK2193874 (80 nM, Fig 2B(ii)) and we have previously shown that the selective Trpv4 activator 4αPDD (1 μM) significantly increased the Po of this Trpv4-like channel on PVN neurones (Feetham, Nunn, Lewis, Dart & Barrett-Jolley, 2015).

**Figure 2.**
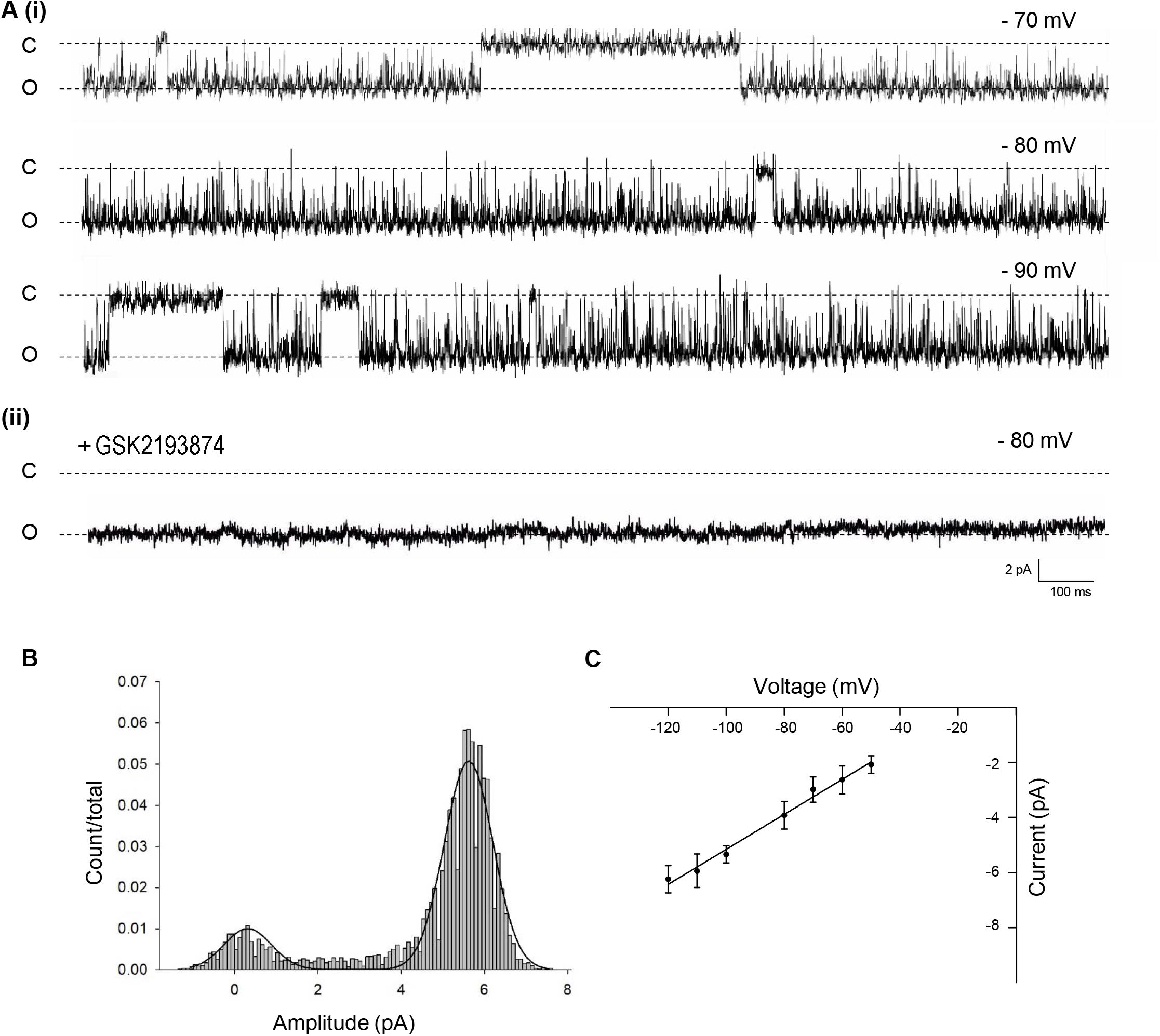
Single channel properties of Trpv4-like channels from PVN neurones. **A.** Representative single-channel current fluctuations through Trpv4-like channels from mouse PVN neurones. Holding potentials are indicated on the trace. The open and closed channel levels are indicated by O and C, respectively. This trace is representative of 10 experiments where Trpv4-like channels were observed and these channels were absent in the presence of the TRPV4 inhibitor GSK2193874. **B.** The amplitude histogram is shown for the trace in A. **C.** Current-voltage relationship for Trpv4-like channels. Mean±SEM is shown (n=9).

### PVN Trpv4-like channels are sensitive to temperature

Decreasing temperature dramatically reduced the Po of this Trpv4-like channel (\**p*< 0.05, \*\*\**p*< 0.001, Fig. 2A and B, n=9). As shown in Fig. 3A, at 37 °C, Trpv4-like channels were predominately open with only brief closing events and at lower temperatures, there was an apparent reduction in open durations. We found that the decrease in mean open probability (Po) was mediated by profound decrease in the mean open time (\*\*\**p*< 0.001, Fig. 3D, n=9) with no change in mean closed time. We also observed a small but significant decrease in Trpv4-like channel conductance when temperature was decreased (Fig. 3B, n=9, blue line).

**Figure 3.**
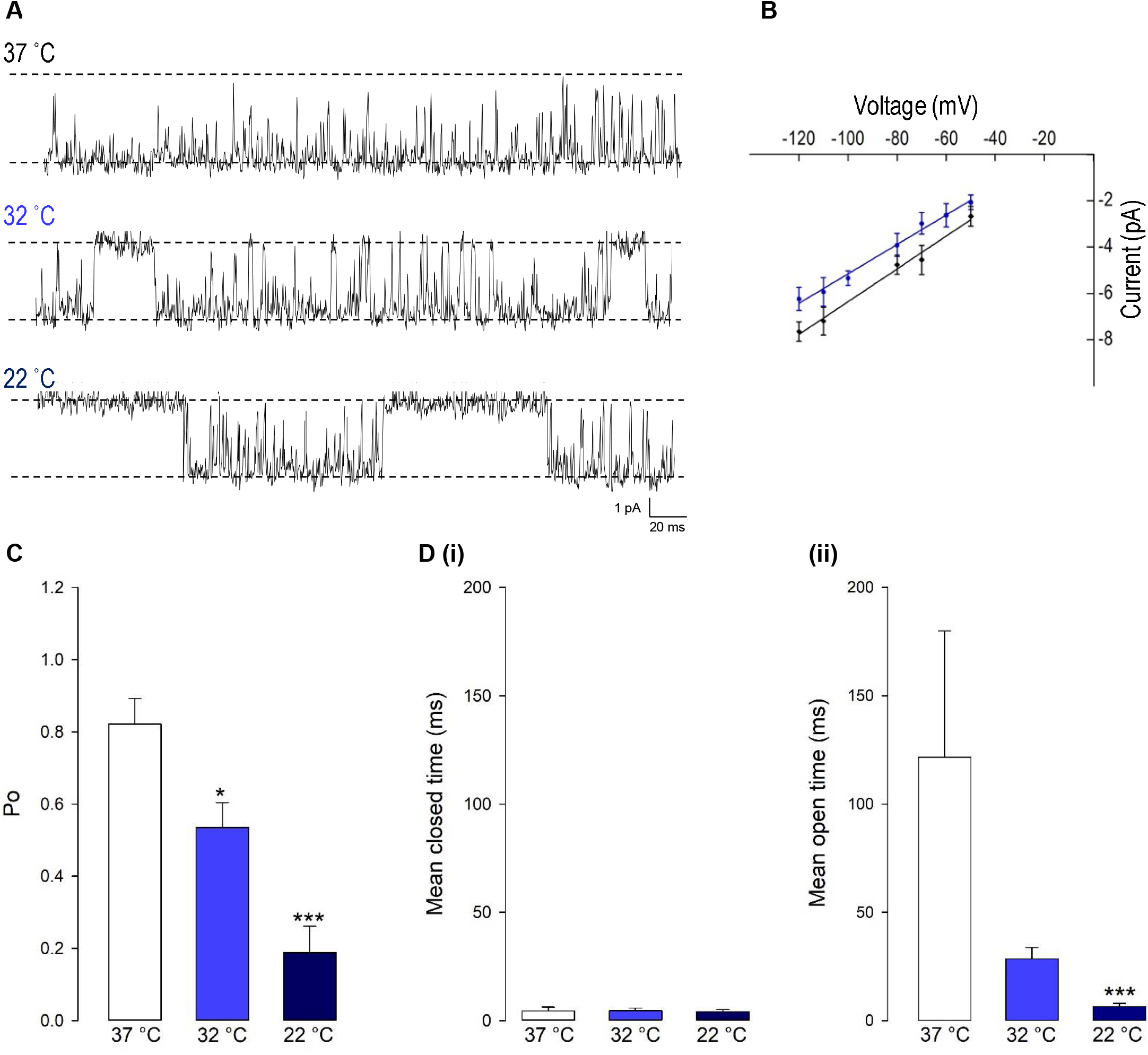
The gating of PVN Trpv4-like channels is temperature sensitive. **A.** Representative single-channel current fluctuations through Trpv4-like channels from mouse PVN neurones at 37°C, 32°C and 22°C. The open and closed channel levels are indicated by O and C, respectively. **B.** Current-voltage relationship for Trpv4-like channels at 37°C (red circles) and 22°C (blue circles). **C.** The *Po* of Trpv4-like channels at 37°C, 32°C and 22°C is shown. D. Mean open and closed times for Trpv4-like channels. Mean±SEM is shown (n=7 at 22°C, n =7 at 32°C and n=9 at 37°C). Where not shown, error bars are within the symbols.

To investigate changes in gating further, we performed detailed analysis of the open and closed dwell-time distributions. The number of time constants (*tau*) was 3 for both the closed and open channel levels (observed in 6/9 experiments). The remaining channels were refit to meet these tau restraints. A representative example is shown in Fig. 4. Lower temperatures markedly reduced 2 out of 3 open *taus* (*tau*_O2_ and *tau*_O3_), with no significant change in any of the closed *taus* (Table 1). Thus, the mechanism for the temperature evoked decrease in Trpv4-like *Po* observed when cooled is a decrease in mean open dwell times.

**Table 1.** Time constants and percentage areas are shown as obtained from maximum likelihood fitting of pdfs to closed and open lifetime distributions of Trpv4-like channels from PVN neurones. Data are presented as mean ± SEM for 7-9 experiments (\**p*<0.05, \*\**p*<0.01, \*\*\**p*<0.0001).

| *Closed* |  |  |  |  |  |
| --- | --- | --- | --- | --- | --- |
| 22 °C |  | 32 °C |  | 37 °C |  |
| tau (ms) | Area (%) | tau (ms) | Area (%) | tau (ms) | Area (%) |
| 0.58±0.01 | 58 | 0.75±0.02 | 65 | 1.24±0.3 | 69 |
| 5.26±0.02 | 30 | 5.87±0.2 | 27 | 7.28±2.3 | 24 |
| 71.22±3.9 | 12 | 49.96±6.9 | 7 | 82.19± | 7 |

| **Open** |  |  |  |  |  |
| --- | --- | --- | --- | --- | --- |
| 22 °C |  | 32 °C |  | 37 °C |  |
| tau (ms) | Area (%) | tau (ms) | Area (%) | tau (ms) | Area (%) |
| 1.32±0.68 | 40 | 1.88±0.41 | 38 | 2.73±0.32 | 28 |
| 3.99±1.65 | 39 | 14.14±2.3** | 39 | 40.07±8.22*** | 49 |
| 41.51±12.06 | 21 | 106.55±19.36* | 23 | 224.57±48.17** | 23 |

**Figure 4.**
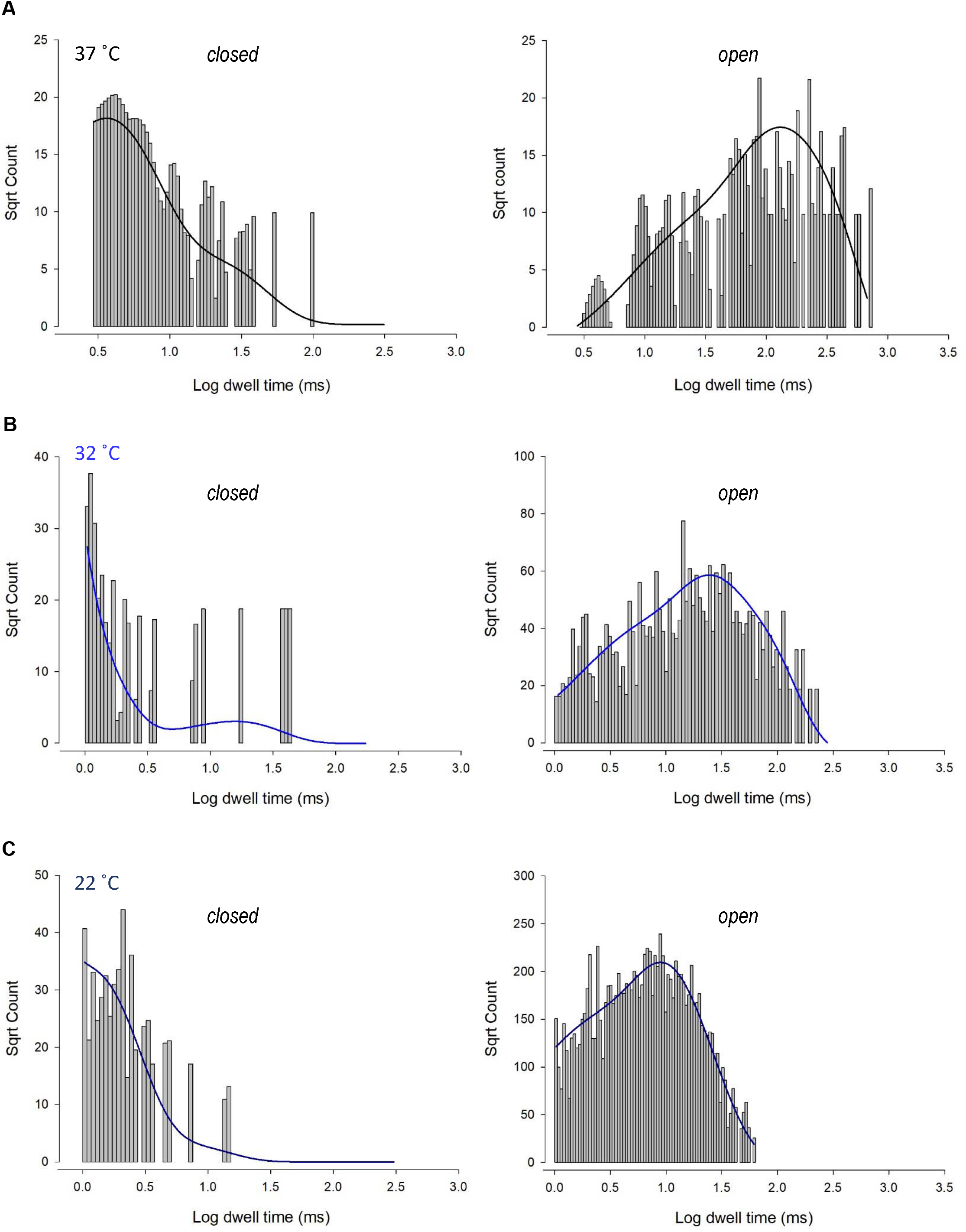
Kinetics of Trpv4-like channels from PVN neurones. Kinetic analysis of Trpv4-like channel dwell-times from PVN neurones recorded in cell attached-patch mode at 37°C (A), 32°C (B) and 22°C (C). Closed (left) and open (right) dwell-times were fitted with 3 exponentials (solid lines). Data are transformed with log-binning (x-axis) and square root of frequency (y-axis) so that exponential time constants are visible as peaks (Sigworth and Sine, 1987). Mean values are given in Table 1 and a kinetic schema in Fig. 5.

### *In silico* analysis of Trpv4 inhibition and prediction of PVN action current frequency

Characterisation of precise single ion channel gating facilitates the computation of neuronal action potential firing properties, which may correlate to how sympathetic output may be controlled. Our working hypothesis of PVN neurones is that as temperature decreases, decreasing activity of calcium permeable TRP channels leads to a decrease in the activity of nearby Ca^2+^-activated potassium channels (K_Ca_), respectively, without causing biologically significant changes in global [Ca^2+^] (Feetham, Nunn, Lewis, Dart & Barrett-Jolley, 2015). We therefore hypothesized that the reduction in Trpv4 activity observed at cooler temperatures would result in an increase in the frequency of spontaneous action currents (ACf) from PVN neurons (Fig. 5B). To test the plausibility of this hypothesis quanitatively, we constructed a mathematical model of a PVN neurone cell based on our previous model (Feetham, Nunn, Lewis, Dart & Barrett-Jolley, 2015), as shown in Fig. 5A. Our stochastic model predicted that decreasing temperature from 37°C to 22°C would increase the frequency of spontaneous action currents (ACf) from PVN neurons (\**p*<0.0001, n=5, Fig. 5C).

**Figure 5.**
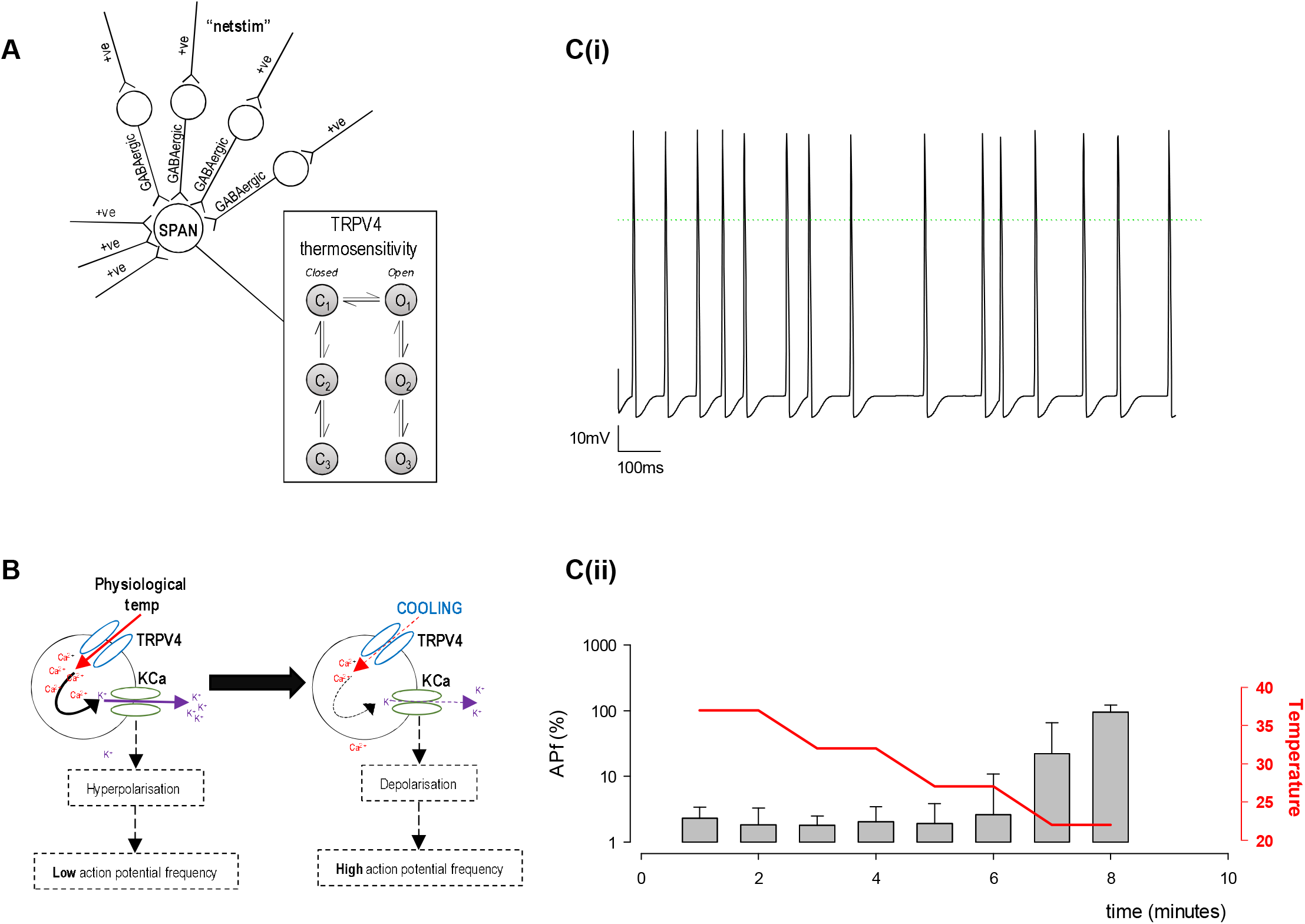
*In silico* model of PVN neurones. **A.** The simple scheme adapted from (Feetham, Nunn, Lewis, Dart & Barrett-Jolley, 2015) whereby influx of Ca^2+^ increases KCa channel activity which hyperpolarizes the cell and increases the inward flux of Ca^2+^, by increasing the driving force for Ca^2+^ entry. **B.** A computer model was adapted from (Feetham, Nunn, Lewis, Dart & Barrett-Jolley, 2015) in NEURON, which includes thermosensitive TRP channels and allows an accumulation of Ca^2+^ into the cell, which is linked to a KCa channel. Within the model, we can change temperature and simulate action currents, shown in **C. D.** Increase in action potential frequency when temperature is decreased (n=5 simulation runs).

### PVN neuronal action current frequency is increased at low temperatures

To validate our mathematical model, we used cell-attached patch-clamp electrophysiology on PVN neurones. We found that ACf was significantly higher at lower temperatures (\*\*\**p*<0.001, Fig. 6, n=7). The maximum effect was observed at room temperature (22°C) where there was a 10-fold increase in ACf (\*\*\**p*<0.001, Fig. 6F, n=7).

**Figure 6.**
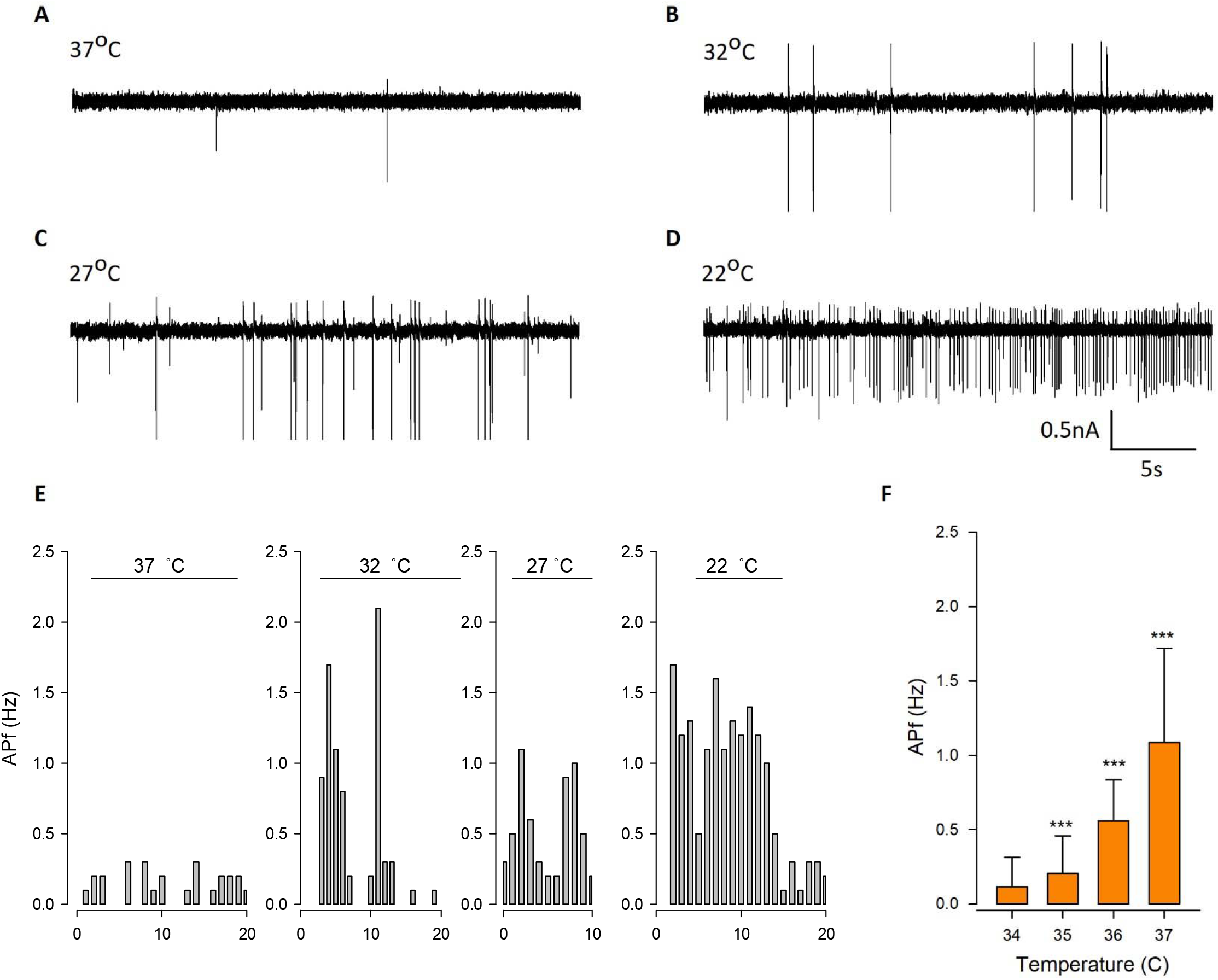
Temperature decreases action current frequency of PVN neurones. **A.** Representative spontaneous firing of action currents from PVN neurones are shown at physiological temperature (37°C) and at lower temperatures of 32°C (B), 27°C (C) and 22°C (D). **E.** Representative frequency histogram showing action current response of a PVN neurone to decreasing temperature. **F.** The mean temperature responses are shown for PVN neurones. Data is presented as mean±SEM (n=6, \*\*\**p*<0.001).

### Temperature sensitivity of PVN neurons

At lower recording temperatures, we observed an increase in ACf which is likely mediated by a decrease in K_Ca_ activity. In this temperature range, a decrease in activity of *any* of our identified warm-activated Ca2+-permeable TRP channels (Trpv4, Trpv3 and Trpm2) could account for this phenomenon (see Fig 1B). We therefore repeated our temperature protocol in the presence of gadolinium (100 μM) which inhibits both Trpv4 (Liedtke et al., 2000) and Trpv3 (Tousova, Vyklicky, Susankova, Benedikt & Vlachova, 2005) and found that while the temperature response persisted from 37°C to 27°C (\*\**p*<0.01, Fig. 7C, n=6) it did not increase further as the temperature was lowered to 22°C. It is worth nothing that at 27°C (the temperature threshold for Trpv4 activation), is where we observed the largest increase in ACf compared to control (\*\**p*<0.01, Fig 6C, n=6).

**Figure 7.**
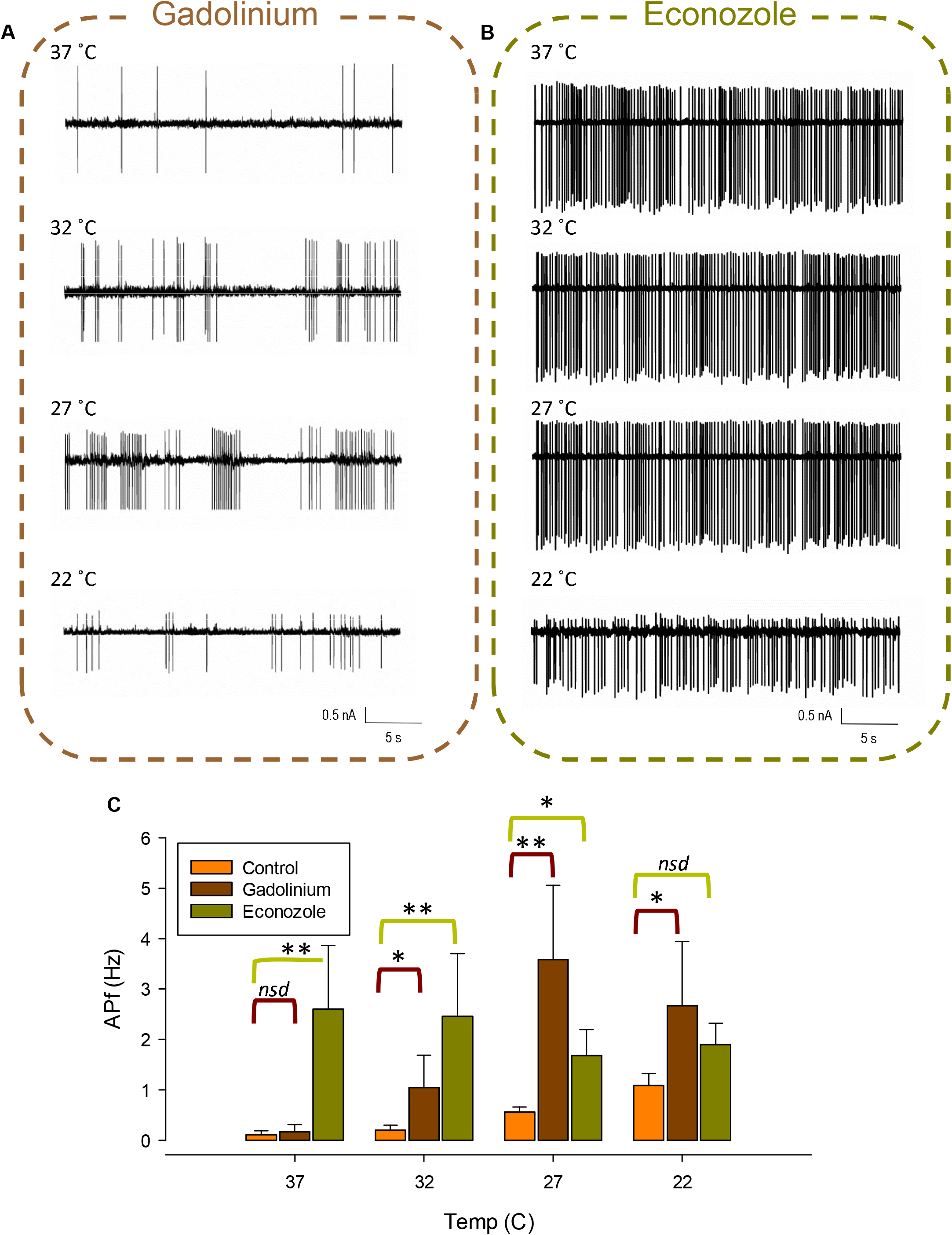
Pharmacological inhibition of various TRP channels on temperature sensitivity of PVN neurones. Representative spontaneous firing of action currents from PVN neurones are shown at physiological temperature (37°C) and at lower temperatures of 32°C, 27°C and 22°C in the presence of (**A**) gadolinium or (**B**) econazole. In **C.**, the mean temperature responses are shown for PVN neurones in the presence of gadolinium (brown) or econozol (green). Data is presented as mean±SEM (n=6 for gadolinium, n=5 for econazole, \**p*<0.05, \*\**p*<0.01).

We also repeated our temperature protocol in the presence of econazole (10 μM) which is known to block the thermosensitive Trpm2 channel (Hill, McNulty & Randall, 2004). We found that ACf no longer changed with recording temperature, indicating that the temperature response was inhibited, (Fig. 7B, n=5). However, at the higher temperatures of 37 °C and 32 °C, ACf was markedly higher to that recorded in control conditions (no drug) (**p<0.01, Fig 7C, n=5).

## Discussion

In this study, we identify thermosensitive TRP channels in the PVN using RT-PCR, of which, Trpv4 is the most abundantly expressed. We characterise the single-channel properties of pharmacologically identified Trpv4-like channels on PVN neurones. We report that these channels are thermosensitive, with decreased activity at lower temperatures, and although our mathematical model predicts that our single channel results could account for the increase in neuronal PVN activity we observed at lower temperatures, we find that the temperature sensitivity of PVN neurones is complex and is likely mediated by the cooperated orchestration of multiple thermosensitive channels, including Trpv4 and Trpm2.

Our results demonstrate that Trpv4-like channels are present on PVN neurones; we have previously illustrated the expression profile of Trpv4 within the PVN using immunohistochemistry (Feetham, Nunn, Lewis, Dart & Barrett-Jolley, 2015) and have shown that application of the selective Trpv4 *agonist* GSK1016790A decreased the firing rate of PVN neurones (Feetham, Nunn, Lewis, Dart & Barrett-Jolley, 2015) and at the level of the whole animal, we have shown that ICV injection of the Trpv4 inhibitor RN1734 prevents the effect of hypotonic ASCF on blood pressure (Feetham, Nunn & Barrett-Jolley, 2015). Here, we characterise the biophysical properties of Trpv4-like channels from PVN neurones using patch-clamp electrophysiology. A Trpv4-like channel was pharmacologically identified with conductance and reversal potential similar to those reported for recombinant Trpv4 channels (Watanabe et al., 2002).

Our results are in agreement with several other studies reporting Trpv4 expression in the PVN; Carreño et al., (2009) show Trpv4-positive cells colocalised with vasopressin (AVP) in both the magnocellular and parvocellular rat PVN (Carreño, Ji & Cunningham, 2009). However, Shenton & Pyner (2018) reported that Trpv4-immunopositive spinally projecting (pre-autonomic) neurone cell bodies were rare in the rat PVN, with immunoreactivity predominately within the magnocellular area (Shenton & Pyner, 2018). This is rather in contrast to our mouse data, both previously (Feetham, Nunn, Lewis, Dart & Barrett-Jolley, 2015) and in this paper where we find Trpv4-like channel activity in 56 % (9/16) of our anatomically and morphologically defined neurones in the parvocellular PVN area. That said, there are major differences between the mouse and rat PVN, for example, unlike the rat PVN, the mouse PVN is not well differentiated and magnocellular and parvocellular neurones are often indistinguishable (Biag et al., 2012). It would now be fascinating to see co-staining of Trpv4 and retrogradely labelled *mouse* PVN too since such data can identify spinally projecting (pre-autonomic) neurones directly. In addition, there have been reported mechanistic species differences too, for example, acute leptin injection induced significant pSTAT3 (a marker of leptin-responsive cells) expression in the rat PVN but not in the mouse. They also reported that the rat PVN exhibited a denser proopiomelanocortin (POMC) innervation compared to the mouse (Campos et al.). In addition, major differences in neuronal populations in different areas of the brain have been reported between species, for example, numerous CRF-ir neurones in the medial preoptic area of rats were barely observed in mice and numerous CRF-ir neurones in the dorsal nucleus of vagus nerve (DMN) of mice that were not present in rats (Wang, Goebel-Stengel, Stengel, Wu, Ohning & Taché, 2011). In addition, Trpv4 channels have been shown to translocate; Baratchi *et al.*, (2016) showed that in human umbilical vein endothelial cells (HUVECs) and in human embryonic kidney 293 cells (HEK293) transfected with Trpv4, sheer stress triggered translocation of Trpv4 to the plasma membrane within seconds of treatment (Baratchi, Almazi, Darby, Tovar-Lopez, Mitchell & McIntyre, 2016). Again in a later publication, they demonstrated that in HUVECs, upon application of shear stress, clusters of Trpv4 channels dispersed into individual channels and translocated from the basolateral to the basal membrane (Baratchi, Knoerzer, Khoshmanesh, Mitchell & McIntyre, 2017). It is therefore plausible that any shear stress or mechanical perturbation may cause additional Trpv4 channels to be translocated to the plasma membrane and be observed more easily at the single channel level.

We find that the gating of Trpv4-like channels is profoundly affected by temperature; *Po* decreased when the temperature was lowered, mediated by a decrease in mean open durations. Trpv4-like channels on PVN neurones were almost maximally activated at normal physiological body temperature, which has been reported elsewhere for TRPV4 channels expressed in HEK293 cells (Watanabe, Vriens, Suh, Benham, Droogmans & Nilius, 2002) and for TRPV4 in isolated hippocampal pyramidal neurons (Shibasaki, Suzuki, Mizuno & Tominaga, 2007). The general consensus is that at physiological temperatures, TRPV4 channels may serve as constitutively open Ca^2+^ entry channels that are sensitive to small deviations in temperature (Watanabe, Vriens, Suh, Benham, Droogmans & Nilius, 2002) and control neuronal excitability *in vitro* and *in vivo* (Shibasaki et al., 2015; Shibasaki, Suzuki, Mizuno & Tominaga, 2007).

Although the molecular dynamics behind heat activation of Trpv4-like channels in the PVN is not known, Watanabe et al., (2002) illustrated that whilst 4αPDD can activate TRPV4 channels in both the cell-attached and cell-free patch clamp configurations, heat application could *only* activate channels in the cell-attached mode, suggesting that there might be an intrinsic heat sensitive ligand or messenger that can active TRPV4 channels from the inside rather than heat activating the channel directly (Watanabe, Vriens, Suh, Benham, Droogmans & Nilius, 2002). We do not know the mechanism behind the heat activation of Trpv4 channels on PVN neurones as we only patched in the cell-attached mode but we could hypothesise that there may be a similar mechanism here.

We have previously shown that pharmacological activation of Trpv4 decreases spontaneous ACf of PVN neurones, mediated by the Ca^2+^-induced activation of K_Ca_ channels (Feetham, Nunn, Lewis, Dart & Barrett-Jolley, 2015). Activation of K_Ca_ channels induces hyperpolarisation, which in turn, draws greater Ca^2+^ into the cell by increasing the driving force for Ca^2+^ entry, setting up a positive feedback loop (Feetham, Nunn, Lewis, Dart & Barrett-Jolley, 2015; Guéguinou, Chantôme, Fromont, Bougnoux, Vandier & Potier-Cartereau, 2014). In our paper and in others, our mathematical model showed that even in the absence of large global changes in Ca^2+^, Trpv4 could permit entry of sufficient Ca^2+^ to activate local SK channels, due to a combinational of local Ca^2+^ signaling domains that limit the diffusion of Ca^2+^ ions after they have entered the cell and the close proximity that often exists between the Ca^2+^ permeable channel (Trpv4 channel) and the Ca^2+^ signaling system (K_Ca_ channel) (Augustine, Santamaria & Tanaka, 2003; Fakler & Adelman, 2008; Neher, 1998)

We have made fundamental changes to our mathematical model. Firstly, we added probabilistic or stochastic gating of ion channels (gating between open and closed states) which attributes to ‘channel noise’ in neuronal activity (Goldwyn & Shea-Brown, 2011; White, Rubinstein & Kay, 2000). It has been demonstrated that the Hodgkin-Huxley derived neuronal models with discrete Markovian ion channel kinetics instead of the usual rate equations can lead to spontaneous generation of action potentials (Hänggi, 2002; Lecar & Nossal, 1971; Skaugen & Walløe, 1979; Strassberg & DeFelice, 1993). In addition, including stochastic behaviour of ion channel gating imparts neuronal noise (Cannon, O’Donnell & Nolan, 2010) that has been shown to effect the variability of spike timing (Schneidman, Freedman & Segev, 1998), firing coherence (Sun, Lei, Perc, Lu & Lv, 2011) and the regularity of spontaneous spike activity (Ozer, Perc & Uzuntarla, 2009). We therefore employed stochastic channels where possible, using the established architecture in NEURON (Hines & Carnevale, 1997b). Also, in the previous model we used intracellular Ca^2+^ buffering that was available in NEURON (Hines & Carnevale, 1997b) but here, we updated to the newer reaction diffusion (RXD) meshwork within pyNeuron (McDougal, Hines & Lytton, 2013a; Newton, McDougal, Hines & Lytton, 2018). This allowed us to model very local changes of Ca^2+^ ions in the direct region of the TRP channel-Ca^2+^-activated potassium channel microdomain. This microdomain approach is critical for understanding functional couplings. Clearly Ca^2+^ concentration does not need to increase across the entire body of the cell, and indeed this is an observation that has been verified in other cell types (Fakler & Adelman, 2008).

Our mathematic simulations predicted that the reduction in Trpv4 *Po* observed at lower temperatures would decrease ACf if the same Trpv4/SK mechanism was at play. In our electrophysiology experiments on neurons in the parvocellular PVN area, we observed a decrease in ACf as we cooled from physiological (37°C) to room (22°C) temperature (Fig. 6). At 37°C, these neurones have little spontaneous activity Fig. 6C), which increased 1.8 fold even with a small 5°C decrease in temperature. Our previous work (Feetham, Nunn, Lewis, Dart & Barrett-Jolley, 2015) was performed at room temperature and therefore in this study, we decreased the temperature of our bath down all the way to 22°C for comparative reasons.

In the presence of gadolinium, which was used to block the warm activated Ca^2+^-permeable TrpvV4 (Liedtke et al., 2000), TRPV3 (Tousova, Vyklicky, Susankova, Benedikt & Vlachova, 2005) and TRPM3 (Kraft et al., 2004) channels, we found that the overall temperature response was largely still present; we observed a decrease in ACf with lower temperatures, however, the effect plateaued after 27°C and we did not see a further decrease when temperature was lowered further to 22°C. We hypothesize that gadolinium is targeting Trpv4 in these experiments; at temperatures of 22°C, 27°C and 32°C, in the presence of gadolinium we see an increase in ACf compared to control, which fits our Trpv4/SK functional coupling model of PVN neurones where a reduction in Trpv4 Po would lead to an increase in ACf. Surprisingly, at 37°C, where we know Trpv4 Po would be very high, we did not see any increase in ACf when Trpv4 was inhibited, indicating that another ion channel may be involved.

The remaining target, Trpm2 has been identified as a heat sensor in the POA (Song et al., 2016), is gadolinium insensitive (Harteneck, 2005) and is activated by temperatures >35 °C (Togashi et al., 2006). We therefore repeated our temperature procedure in the presence of 10 μM econazole which is known to inhibit Trpm2 currents (Hill, McNulty & Randall, 2004) and found that the temperature effect on ACf at all temperatures was blocked, indicating a role for Trpm2 in thermosensing in the PVN. In the presence of econazole, compared to control recordings (no drug), we found that ACf was increased at all temperatures apart from room temperature (22°C) and the effect was most pronounced at 37°C which is just above is the temperature activation threshold of Trpm2.

Interestingly, Song *et al.*, (2016) suggested that Trpm2 in preoptic hypothalamic neurones modulates fever responses via the PVN (Song et al., 2016). They found that inhibition of TRPM2^+^ POA neurones resulted in a significant increase in T_c_. This may fit in with our results; we did not patch isolated PVN neurones, but PVN neurones in their somewhat native neuronal environment and thus, any interference from neighboring hypothalamic nuclei such as the POA may be preserved in our brain slice experiments. We show that at 37°C, inhibition of Trpm2 results in a 10-fold increase in ACf, which we hypothesize would lead to an increased sympathetic output and vasoconstriction, which may result in an increase in T_c_. In our single channel electrophysiology experiments, we did not observe a TRPM2-like channel, so we cannot comment on the presence of Trpm2 on PVN neurones or add to the kinetic profile of Trpm2.

We propose that both Trpv4 and Trpm2 are necessary in combination to account for our electrophysiological results; at physiological temperatures we know Trpv4 activity will be high (Fig. 3A) and we assume that Trpm2 will be active as we are over the temperature activation threshold. Inhibiting Trpv4 (with gadolinium) had no effect on ACf, presumably because TRPM2 is still active and conducting enough Ca^2+^ to maintain SK channel activity.

This may be surprising considering Trpv4 has a permeability ratio Ca^2+^ to Na^+^ (PCa/Pna) of 6 (Clapham, Montell, Schultz & Julius, 2003), whereas Trpm2 PCa/PNa is approximately only 0.7 (Kraft et al., 2004; Sano et al., 2001) but we know from our mathematical model that only a small about of Ca^2+^ entry is required to activate nearby K_Ca_ channels. Further work is necessary to confirm the presence of TRPM2 (as opposed to a closely related channel) in the PVN and to determine the comparative expression profile of Trpv4 and Trpv2. As we cool to room temperature, we presume TRPM2 be switched off first (sub 35°C) whereas Trpv4 will be constitutively active even at room temperature, albeit with low Po (Fig. 3A). Inhibiting Trpm2 at room temperature has no significant effect on ACf (presumably as Trpm2 is already switched off), whereas inhibition of Trpv4 results in a significant increase in ACf, suggesting that Trpv4 plays a role in determining neuronal activity at 22°C. We find that blocking Trpm2 with econozole appears to inhibit the temperature effect; ACf is markedly increased (compared to control) at 37 °C and remains consistently high as the temperature is lowered. In addition, in our mathematical model, both the presence of Trpv4 and Trpm2 were necessary to account for our physiological data, further suggesting that multiple TRP channels orchestrate the observed response.

In conclusion, our data suggest that cooling temperature challenge inhibits multiple TRP channels including Trpv4 and Trpm2. Our mathematical model predicts that resulting decreases in intracellular Ca^2+^ would inhibit local SK channels, depolarise neurones and hence increase ACf and our experimental patch-clamp data validates this. Together, these data give insight into the important fundamental mechanisms by which the body decodes temperature signals and maintains homeostasis in an area of the brain adapted to control of the cardiovascular control system.

Table 2? Markov model tables?

## Supplementary figures

**Supplementary Figure 1.**
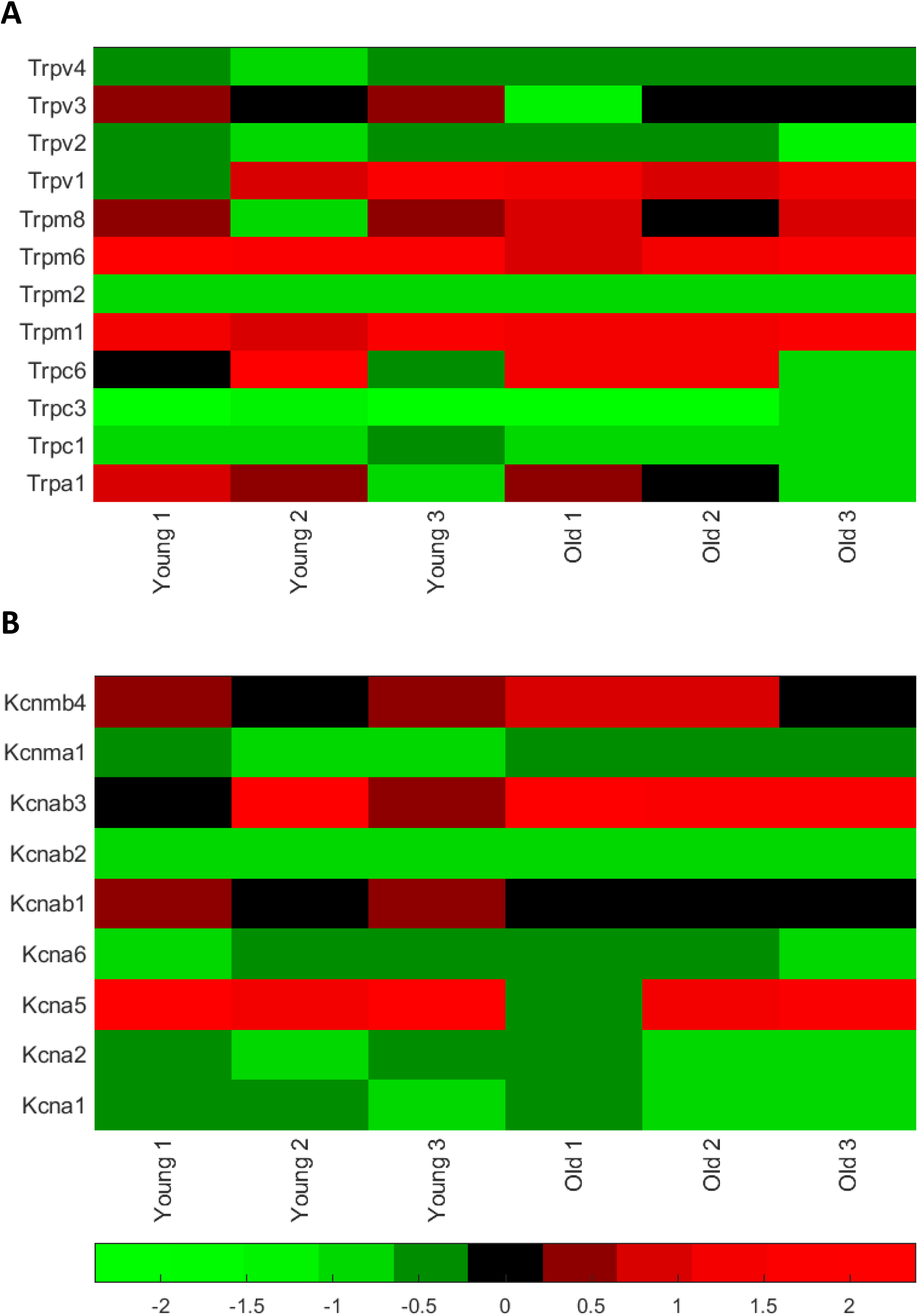
Ion channel gene expression in PVN punches from mice; TRP channels and calcium-activated potassium channels. Standardized DCt levels for mRNA in punches of the PVN from 3 young (6-8 month) and 3 old (26 month) animals. (A) 12 different TRP channel genes. (B) 9 calcium-activated potassium channel related genes. The colour bar pertains to A and B. Since this DCt, the green (negative values) are relatively high expression and the red (positive values) are low expression. Raw data in Supplementary Table 1.

**Supplementary Figure 2.**
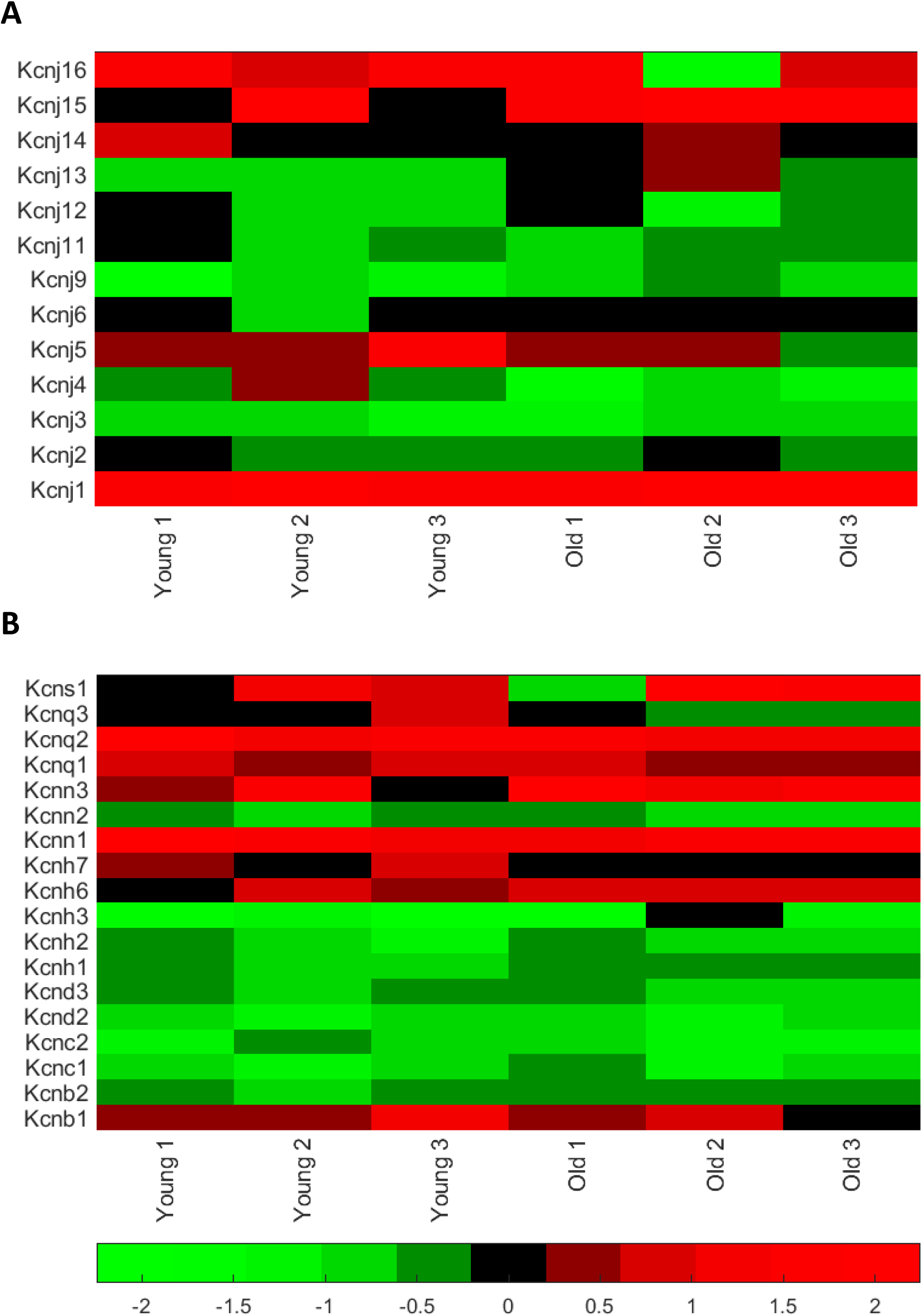
Ion channel gene expression in PVN punches from young and old mice; inwardly rectifying potassium channels and other potassium channels. Standardized DCt levels for mRNA in punches of the PVN from 3 young (6-8 month) and 3 old (26 month) animals. (A) Inwardly rectifying potassium channel genes. (B) Other potassium channel related genes. The colour bar pertains to A and B. Since this DCt, the green (negative values) are relatively high expression and the red (positive values) are low expression. Raw data in Supplementary Table 1.

**Supplementary Figure 3.**
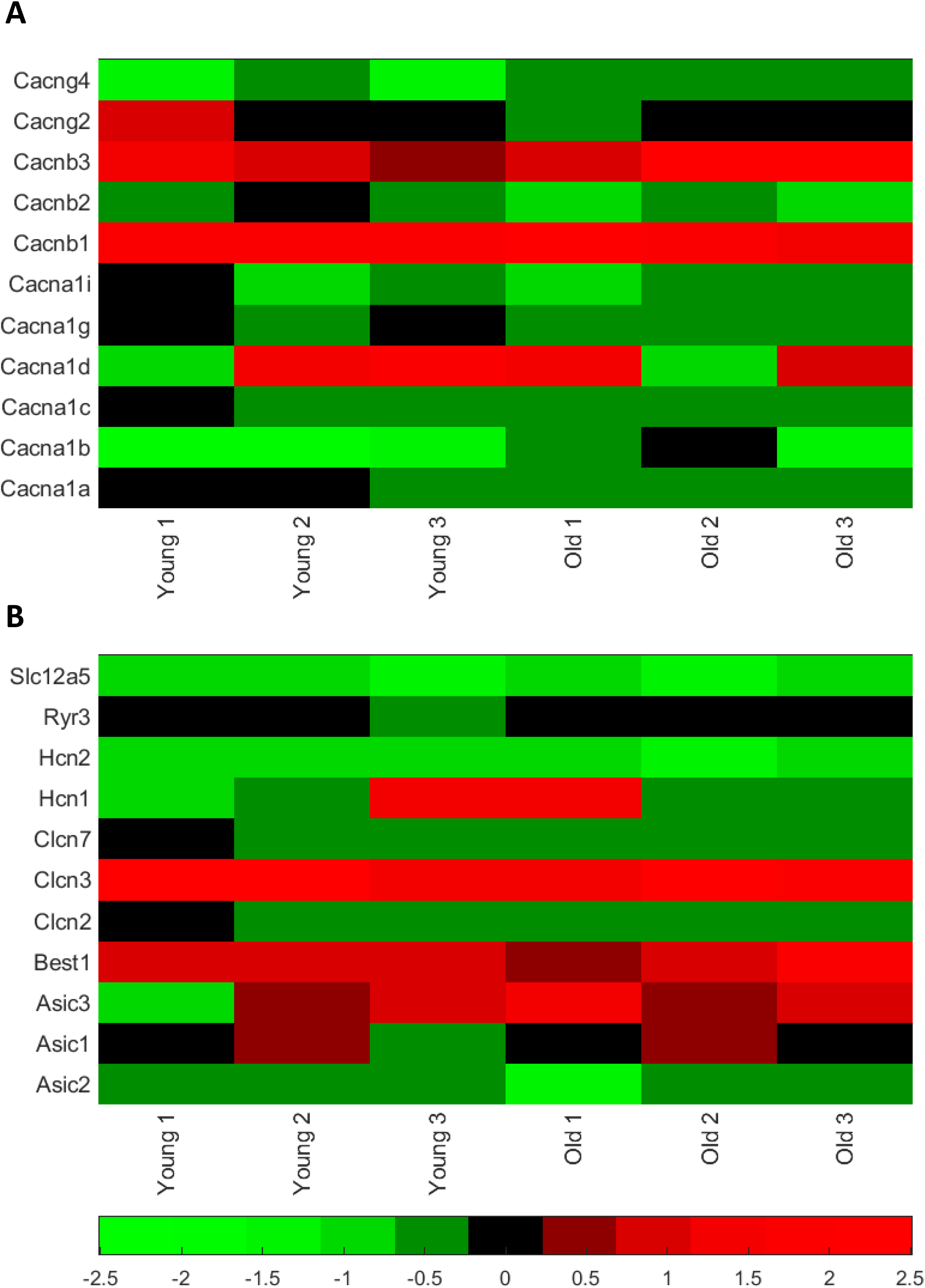
Ion channel gene expression in PVN punches from young and old mice; calcium ion channels and other ion channels. Standardized DCt levels for mRNA in punches of the PVN from 3 young (6-8 month) and 3 old (26 month) animals. (A) Calcium voltage-gated ion channel related genes. (B) Other ion channel related genes, not included in the above families together with the transporter Slc12a5. The colour bar pertains to A and B. Since this DCt, the green (negative values) are relatively high expression and the red (positive values) are low expression. Raw data in Supplementary Table 1.

**Supplementary Table 1:**
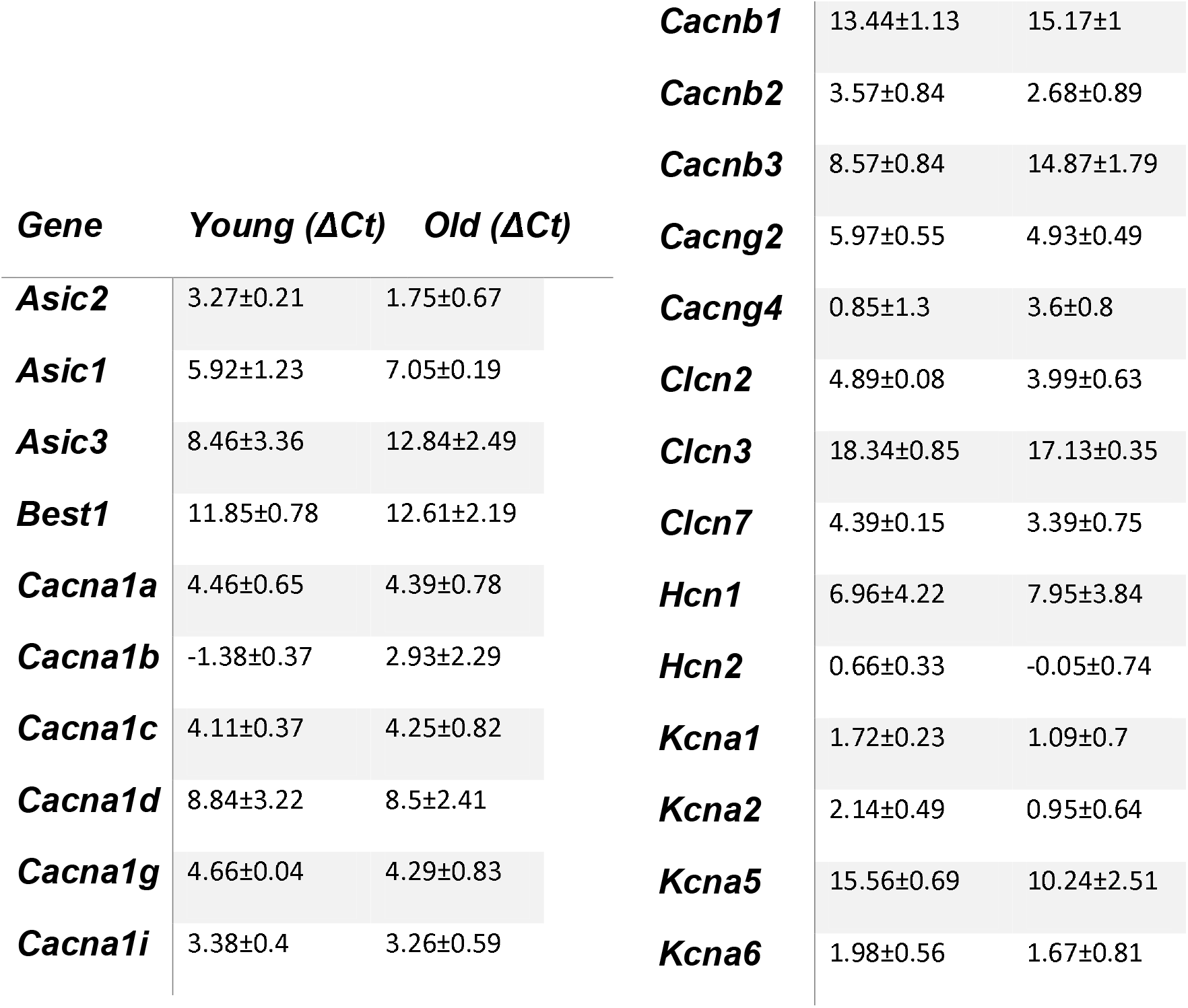
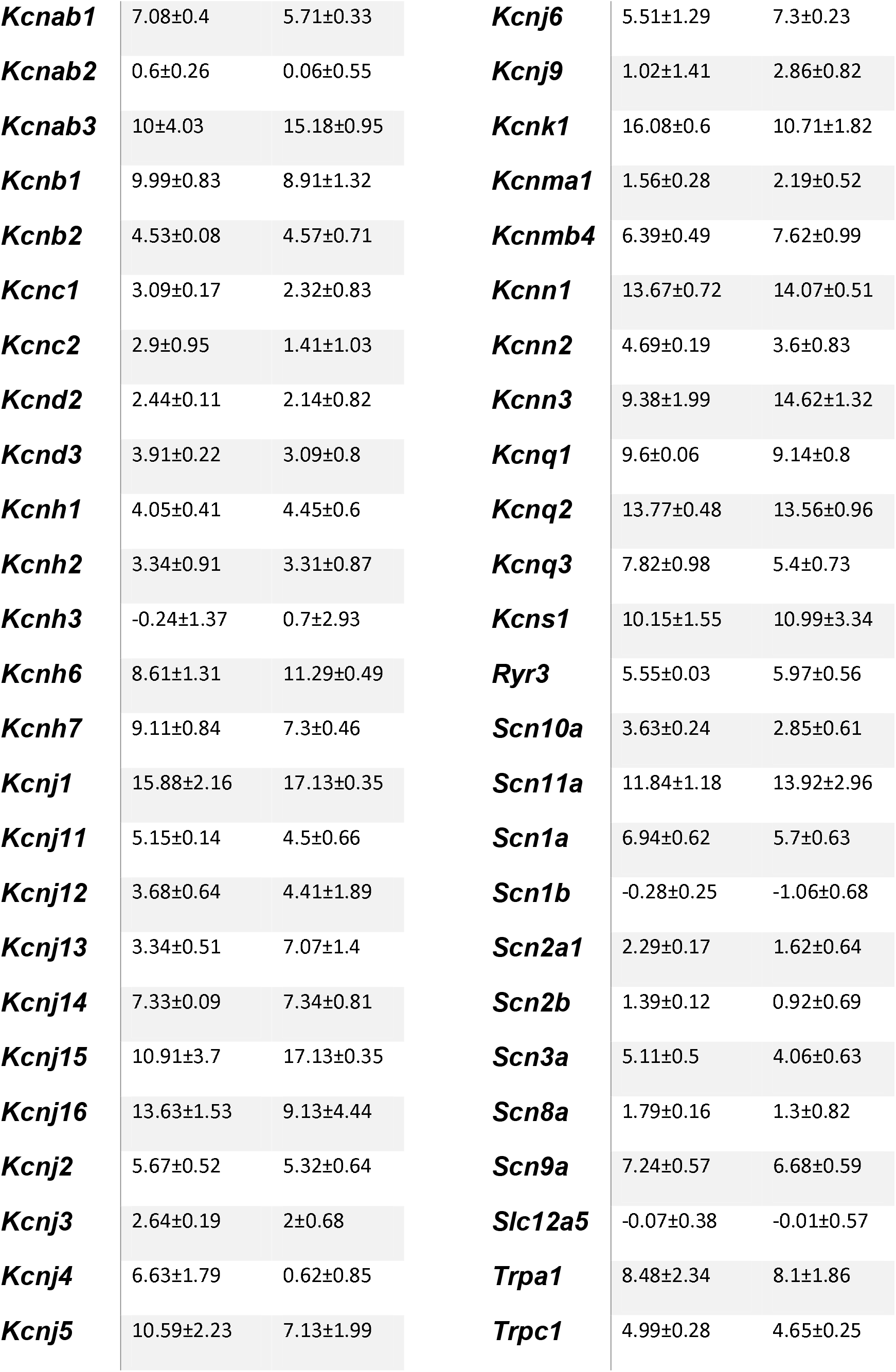
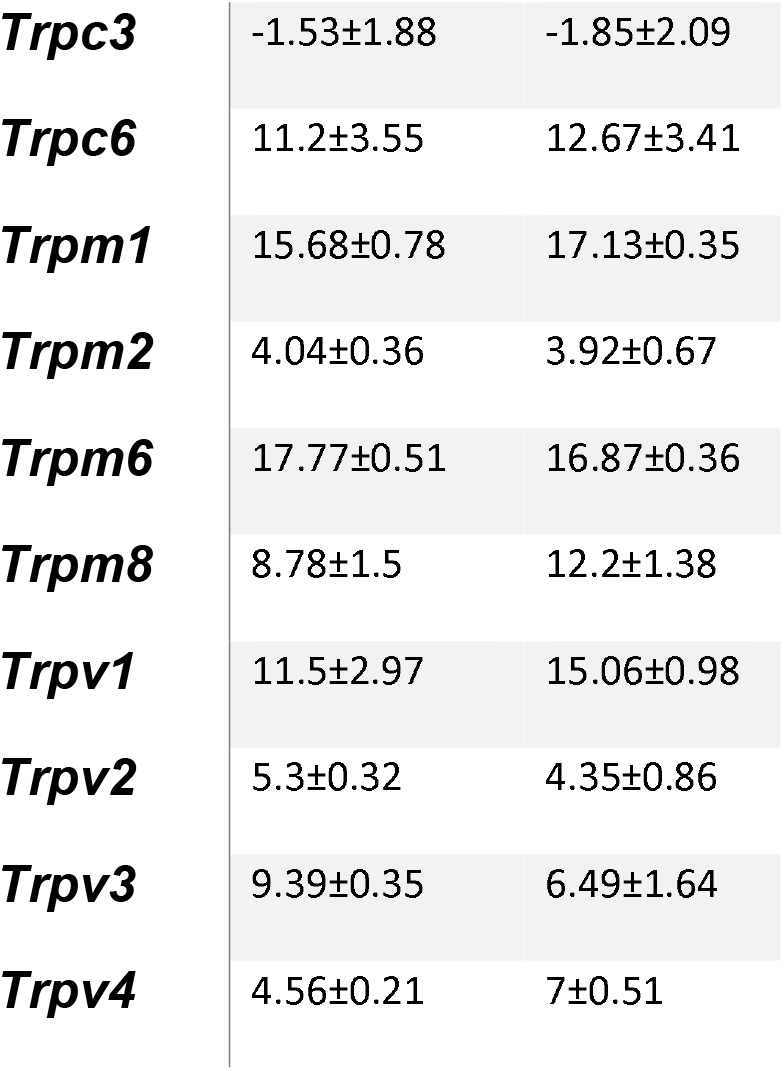
Expression relative to mean of 3 housekeeper gene (Actb, Ldha, Rplp1) Cts for that sample is presented as the ΔCt. *n*=3 for each value.

| Gene | Young ($\Delta Ct$) | Old ($\Delta Ct$) |
| --- | --- | --- |
| **Asic2** | 3.27±0.21 | 1.75±0.67 |
| **Asic1** | 5.92±1.23 | 7.05±0.19 |
| **Asic3** | 8.46±3.36 | 12.84±2.49 |
| **Best1** | 11.85±0.78 | 12.61±2.19 |
| **Cacna1a** | 4.46±0.65 | 4.39±0.78 |
| **Cacna1b** | -1.38±0.37 | 2.93±2.29 |
| **Cacna1c** | 4.11±0.37 | 4.25±0.82 |
| **Cacna1d** | 8.84±3.22 | 8.5±2.41 |
| **Cacna1g** | 4.66±0.04 | 4.29±0.83 |
| **Cacna1i** | 3.38±0.4 | 3.26±0.59 |
| **Cacnb1** | 13.44±1.13 | 15.17±1 |
| **Cacnb2** | 3.57±0.84 | 2.68±0.89 |
| **Cacnb3** | 8.57±0.84 | 14.87±1.79 |
| **Cacng2** | 5.97±0.55 | 4.93±0.49 |
| **Cacng4** | 0.85±1.3 | 3.6±0.8 |
| **Clcn2** | 4.89±0.08 | 3.99±0.63 |
| **Clcn3** | 18.34±0.85 | 17.13±0.35 |
| **Clcn7** | 4.39±0.15 | 3.39±0.75 |
| **Hcn1** | 6.96±4.22 | 7.95±3.84 |
| **Hcn2** | 0.66±0.33 | -0.05±0.74 |
| **Kcna1** | 1.72±0.23 | 1.09±0.7 |
| **Kcna2** | 2.14±0.49 | 0.95±0.64 |
| **Kcna5** | 15.56±0.69 | 10.24±2.51 |
| **Kcna6** | 1.98±0.56 | 1.67±0.81 |
| ***Kcnab1*** | 7.08±0.4 | 5.71±0.33 |
| ***Kcnab2*** | 0.6±0.26 | 0.06±0.55 |
| ***Kcnab3*** | 10±4.03 | 15.18±0.95 |
| ***Kcnb1*** | 9.99±0.83 | 8.91±1.32 |
| ***Kcnb2*** | 4.53±0.08 | 4.57±0.71 |
| ***Kcnc1*** | 3.09±0.17 | 2.32±0.83 |
| ***Kcnc2*** | 2.9±0.95 | 1.41±1.03 |
| ***Kcnd2*** | 2.44±0.11 | 2.14±0.82 |
| ***Kcnd3*** | 3.91±0.22 | 3.09±0.8 |
| ***Kcnh1*** | 4.05±0.41 | 4.45±0.6 |
| ***Kcnh2*** | 3.34±0.91 | 3.31±0.87 |
| ***Kcnh3*** | -0.24±1.37 | 0.7±2.93 |
| ***Kcnh6*** | 8.61±1.31 | 11.29±0.49 |
| ***Kcnh7*** | 9.11±0.84 | 7.3±0.46 |
| ***Kcnj1*** | 15.88±2.16 | 17.13±0.35 |
| ***Kcnj11*** | 5.15±0.14 | 4.5±0.66 |
| ***Kcnj12*** | 3.68±0.64 | 4.41±1.89 |
| ***Kcnj13*** | 3.34±0.51 | 7.07±1.4 |
| ***Kcnj14*** | 7.33±0.09 | 7.34±0.81 |
| ***Kcnj15*** | 10.91±3.7 | 17.13±0.35 |
| ***Kcnj16*** | 13.63±1.53 | 9.13±4.44 |
| ***Kcnj2*** | 5.67±0.52 | 5.32±0.64 |
| ***Kcnj3*** | 2.64±0.19 | 2±0.68 |
| ***Kcnj4*** | 6.63±1.79 | 0.62±0.85 |
| ***Kcnj5*** | 10.59±2.23 | 7.13±1.99 |
| ***Kcnj6*** | 5.51±1.29 | 7.3±0.23 |
| ***Kcnj9*** | 1.02±1.41 | 2.86±0.82 |
| ***Kcnk1*** | 16.08±0.6 | 10.71±1.82 |
| ***Kcnma1*** | 1.56±0.28 | 2.19±0.52 |
| ***Kcnmb4*** | 6.39±0.49 | 7.62±0.99 |
| ***Kcnn1*** | 13.67±0.72 | 14.07±0.51 |
| ***Kcnn2*** | 4.69±0.19 | 3.6±0.83 |
| ***Kcnn3*** | 9.38±1.99 | 14.62±1.32 |
| ***Kcnq1*** | 9.6±0.06 | 9.14±0.8 |
| ***Kcnq2*** | 13.77±0.48 | 13.56±0.96 |
| ***Kcnq3*** | 7.82±0.98 | 5.4±0.73 |
| ***Kcns1*** | 10.15±1.55 | 10.99±3.34 |
| ***Ryr3*** | 5.55±0.03 | 5.97±0.56 |
| ***Scn10a*** | 3.63±0.24 | 2.85±0.61 |
| ***Scn11a*** | 11.84±1.18 | 13.92±2.96 |
| ***Scn1a*** | 6.94±0.62 | 5.7±0.63 |
| ***Scn1b*** | -0.28±0.25 | -1.06±0.68 |
| ***Scn2a1*** | 2.29±0.17 | 1.62±0.64 |
| ***Scn2b*** | 1.39±0.12 | 0.92±0.69 |
| ***Scn3a*** | 5.11±0.5 | 4.06±0.63 |
| ***Scn8a*** | 1.79±0.16 | 1.3±0.82 |
| ***Scn9a*** | 7.24±0.57 | 6.68±0.59 |
| ***Slc12a5*** | -0.07±0.38 | -0.01±0.57 |
| ***Trpa1*** | 8.48±2.34 | 8.1±1.86 |
| ***Trpc1*** | 4.99±0.28 | 4.65±0.25 |
| ***Trpc3*** | -1.53±1.88 | -1.85±2.09 |
| ***Trpc6*** | 11.2±3.55 | 12.67±3.41 |
| ***Trpm1*** | 15.68±0.78 | 17.13±0.35 |
| ***Trpm2*** | 4.04±0.36 | 3.92±0.67 |
| ***Trpm6*** | 17.77±0.51 | 16.87±0.36 |
| ***Trpm8*** | 8.78±1.5 | 12.2±1.38 |
| ***Trpv1*** | 11.5±2.97 | 15.06±0.98 |
| ***Trpv2*** | 5.3±0.32 | 4.35±0.86 |
| ***Trpv3*** | 9.39±0.35 | 6.49±1.64 |
| ***Trpv4*** | 4.56±0.21 | 7±0.51 |

